# Stem cell lineage survival as a noisy competition for niche access

**DOI:** 10.1101/2020.02.12.945253

**Authors:** Bernat Corominas-Murtra, Colinda L.G.J. Scheele, Kasumi Kishi, Saskia I.J. Ellenbroek, Benjamin D. Simons, Jacco van Rheenen, Edouard Hannezo

## Abstract

Understanding to what extent stem cell potential is a cell-intrinsic property, or an emergent behavior coming from global tissue dynamics and geometry, is a key outstanding question of stem cell biology. Here, we propose a theory of stem cell dynamics as a stochastic competition for access to a spatially-localized niche, giving rise to a “stochastic conveyor-belt” model. Cell divisions produce a steady cellular stream which advects cells away from the niche, while random rearrangements enable cells away from the niche to be favourably repositioned. Importantly, even when assuming that all cells in a tissue molecularly equivalent, the model predicts a common (“universal”) functional dependence of the long-term clonal survival probability on the position within the niche, as well as the emergence of a well-defined number of “functional” stem cells, dependent only on the rate of random movements vs. mitosis-driven advection. We test the predictions of this theory on datasets on pubertal mammary gland tips, embryonic kidney tips as well homeostatic intestinal crypt, and find good quantitative agreement for the number of functional stem cells in each organ, as well as the predicted functional dependence of the competition.

---

Many biological tissues are renewed via small numbers of stem cells, which divide to produce a steady stream of differentiated cells and balance homeostatic cell loss. Although novel experimental approaches in the past decade have produced key insights into the number, identity, and (often stochastic) dynamics of stem cells in multiple organs, an outstanding question remains as to whether stem cell potential is a cell-intrinsic, “inherited” property, or rather an extrinsic, context-dependent state emerging from the collective dynamics of a tissue and cues from local “niches”, or microenvironments [1–8]. Although recent experiments have provided evidence for the latter in settings such as the growing mammary gland [9], adult interfollicular epidermis [10, 11], spermatogenesis [12] or the intestinal epithelium [13], a more global theoretical framework allowing to quantitatively interpret these findings is still lacking.

The case of the intestinal crypt serves as a paradigmatic example of the dynamics of tissue renewal, and is one of the fastest in mammals [13]. The intestinal crypt consists of a small invagination in the intestine where the epithelial cells populating the intestinal walls are constantly produced. The very bottom of the crypt hosts a small number of proliferative, Lgr5+ stem cells, [14] that divide and push the cells located above them to the transit amplification (TA) region, where cells lose self-renewal potential. Cells are eventually shed in the villus a few days later, constituting a permanent “conveyor-belt” dynamics. Lineage tracing approaches, which irreversibly label a cell and its progeny [3], have been used to ask which cell type will give rise to lineages that renew the whole tissue and have revealed that all Lgr5+ cells can stochastically compete in an equipotent manner on the long term [15–18], but still display positional-dependent short term biases for survival [13]. Interestingly, similar conclusions have been achieved in pubertal mammary gland development [9], where branching morphogenesis is performed through the proliferation of the cells in the terminal end buds of the ducts, the region where the mammary stem cells (MaSCs) reside [9, 19]. In both cases, intravital imaging revealed random cellular motions enabling a cell to move against the cellular flow/drift defined by the conveyor belt dynamics. Moreover, in the intestine, tissue damage, or genetic ablation of all Lgr5+ stem cells, caused Lgr5-cells to recolonize the crypts and re-express Lgr5+ to function as stem cells [13], arguing for extensive reversibility and flexibility in the system [20]. In addition, Lgr5- and Lgr5+ cells were also shown to nearly equally contribute to intestinal morphogenesis [21]. Altogether, this supports proposals that the definition of stem cell potential should evolve to emphasize, instead of molecular markers, the functional ability of replace lost cells via mitosis [22, 23] and survive long-term in tissues. However, this new definition raises a number of outstanding conceptual problems: What then defines the number of functional stem cells in a tissue? How can short-term biases be reconciled with long-term equipotency? Is there a sharp distinction between stem and non-stem cells, or is there instead a continuum of stem cell potential together with flexible transition between states? Qualitatively, it is clear that fluctuations and positional exchange are needed to prevent a single cell in the most favourable position to be the unique “functional” stem cell (defined as cells whose lineage colonizes a tissue compartment on the long-term). Incorporating these features in a dynamical model of stem cell growth and replacement, able to make predictions e.g., over the probability of lineage perpetuation, would represent an important step towards the understanding of how stem cells operate in the process of tissue development and renewal.

In this paper we develop a reaction-diffusion formalism for stem cell renewal in the presence of noise and local niches, taking into account local tissue geometry as well as cell division and random cell movements –see Fig. (1a–c). Importantly, within this purely extrinsic and dynamical approach, which does not need to posit any intrinsic “stem cell identity”, a well-defined number of functional stem cell emerges, which only depends on the geometry and a balance between the noisiness of cell movements and division rates advecting cells away from niche regions. This model also predicts that “stem cell potential” should decay continuously as a function of distance from the niche, with a “universal” Gaussian functional dependence. We test this prediction against published live-imaging datasets for the homeostatic intestinal crypt [13] and during the branching of embryonic kidney explants [24], and find a good quantitative agreement for the full survival probability of cells depending on their initial position relative to the niche. Furthermore, we use our theoretical results to extract the amplitude of the random positional fluctuations in the developing mammary gland using lineage tracking experiments [9]. This enables us to predict the number of functional stem cells for this system, finding values fully consistent with estimates previously reported.

**FIG. 1:**
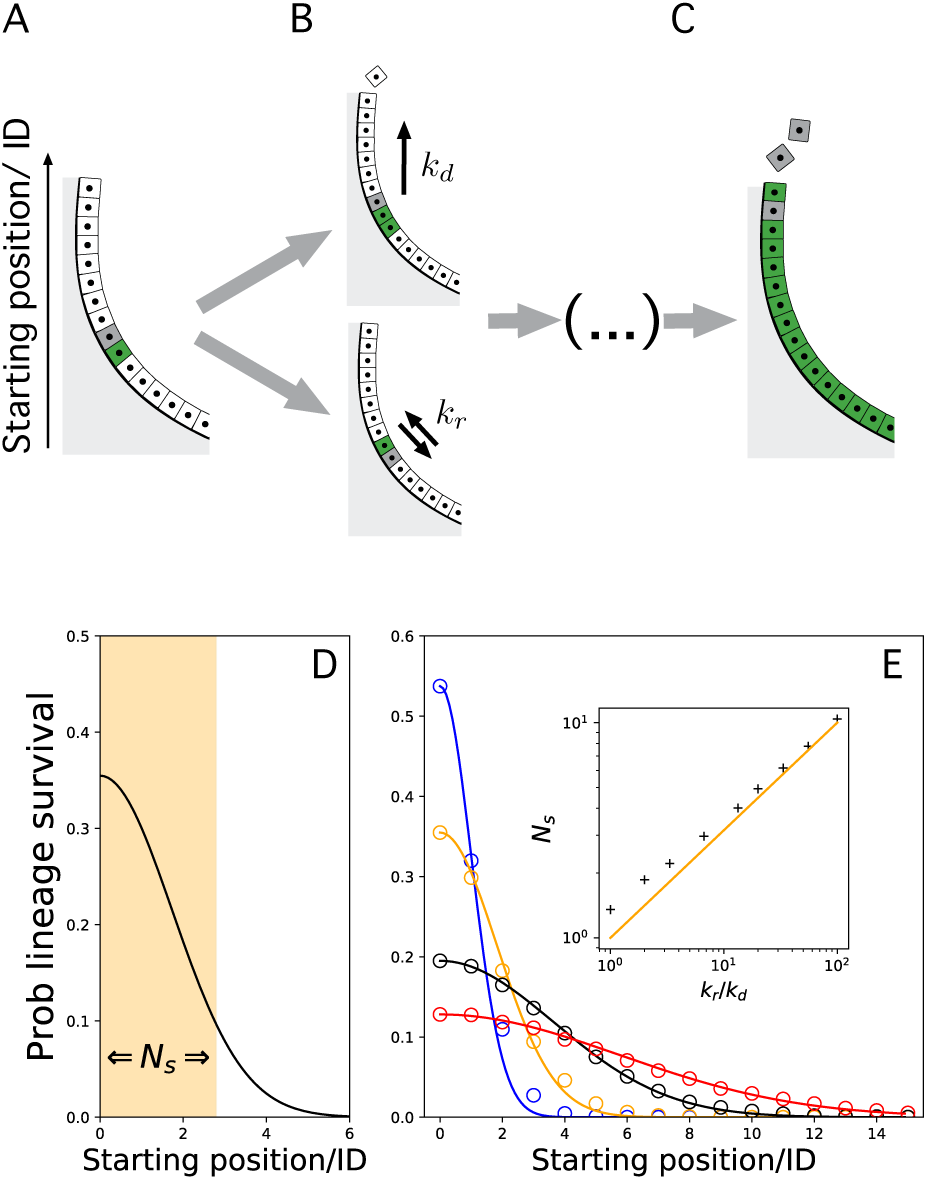
Stochastic conveyor belt as a paradigm for stem cell renewal. A/ A cell in the epithelial wall of the crypt can B/ duplicate at rate *k*_*d*_ pushing the upper cells up, creating a conveyor-belt mechanism; or switch its position randomly at rate *k*_*r*_, introducing a stochastic or noisy ingredient in the dynamics. C/ At longer time scales, the lineage of a single starting cell colonizes the whole system. D/ The probability that a given lineage colonizes the entire system depends critically on the initial position of its mother cell with respect to the bottom of the system. The probability decays as a Gaussian with the distance from the origin (black solid line), see text for details. The functional number of stem cells emerges from the dynamics as the amount cells by which fluctuations may result in a high probability of colonizing the entire system. Since the amplitude of fluctuations is 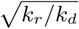, the cells located in the interval 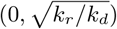 define the functional stem cell region, highlighted in orange, and referred as *N*_*s*_ (see text). Here we plot the probability of lineage survival for *k*_*r*_*/k*_*d*_ = 3. E/ Long term survival probability *p*(*c*_*n*_) as a function of initial starting position *n* = 0, 1, 2, …, with respect to the base of the system for several values of *k*_*r*_*/k*_*d*_ (1, 3.3, 13.3 and 33 in resp. blue, orange, black and red). Dots show simulation results and lines show the analytical prediction 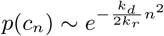, as shown in equation (3). In the inset, analytical criteria with best-fit to the variance (black crosses) against the prediction 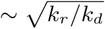 (orange solid line).

## I. DYNAMICS OF TISSUE RENEWAL AND DEVELOPMENT

To develop the model, we first consider the simplest situation of a one-dimensional column of cells, with a rigid boundary condition at the base (mimicking, for instance, the bottom of the crypt), so that each cell division produces a pushing force upwards transmitted to the cells above (or in the case of growing mammary gland or kidney, driving ductal elongation). This model is motivated by its simplicity, as it is able to qualitatively derive the essential traits of the complex dynamics studied here. As we shall see, further refinements, aimed at making predictions for real systems, consider much more realistic geometries. From this simple dynamics, we define the number of functional stem cells as the typical number of cells that have a non-negligible probability to produce long-term progenies (without “losing” the competition against other cells). If the dynamics was fully devoid of noise (a simple conveyor belt) and all cell divisions were symmetric, then one of the bottom-most cells would always win the competition. Thus, there would be a single row of functional stem cells, which is the limiting case of described in Ref. [16] of symmetric and stochastic 1D competition along a ring of equipotent cells. However, as mentioned above, an extensive amount of noise in cellular movements and rearrangement is observed, to different levels, in multiple settings via live-imaging, for instance in the mammary gland [9], kidney morphogenesis [24, 25], or intestinal crypts [13]. Intuitively, such rearrangements are expected to increase the number of “functional” stem cells, as re-arrangements allow cells away from the niche to relocate in favourable positions, and would thus provide a biophysical mechanism for setting the number of stem cells assumed in models such as Ref. [16].

### A. Lineage dynamics

To further develop our intuition, and provide a quantitative criterion for how re-arrangements affect the number of cells which participate effectively in the competition, we defined a minimal model of such “stochastic-conveyor belt” dynamics. We start with the equation that describes the stochastic movement of a single cell: First, one must account for the push-up force due to the proliferation of cells located lower in the system at rate *k*_*d*_. Furthermore, it must incorporate a probability for cells to randomly switch position, accounting for the random positional fluctuations, with amplitude *k*_*r*_ (although the exact nature and implementation of this noise does not impact the results, as shown and discussed in Appendix, sections A1 and C). With these ingredients, the stochastic dynamics is governed by the equation:

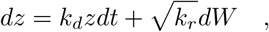

where *dW* is the standard Brownian differential and *z* the distance of the cell from the origin. This dynamics represents an *Ornstein-Uhlenbeck process* [26, 27], with the peculiarity that the drift term is positive. To analyze the evolution of the lineage, we have to introduce a proliferative term in the dynamics. Let *ρ*_*n*_(*z, t*) be the expected density of the lineage *n* at time *t* at position *z*, where *n* is defined such that the mother cell occupies position *z* = *n* at *t* = 0. At the next time step, *k*_*d*_*ρ*_*n*_(*z, t*) new cells of this lineage will appear at position *z* due to cell proliferation. The lineage density evolution dynamics is thus described by a reaction-diffusion equation [28, 29], in which the reaction term describes the exponential growth of the lineage density and the diffusion term follows an Ornstein-Uhlenbeck-like dynamics (see Fig. (1a,b) and the Appendix, sections A1,2):

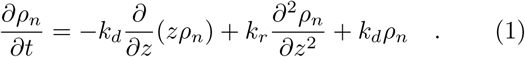

Note that here we have assumed that all cells are of the same type and that properties such as cell divisions and rearrangements do not change spatially; that is, *k*_*r*_ and *k*_*d*_ are constant throughout the system, two assumptions that we lift in subsequent sections. This particular stochastic process has no stationary solutions. However, considering initial conditions *t*_0_ = 0, *z*_0_ = *δ*(*z* − *n*) and natural boundary conditions, one can still compute the time-dependent solutions, which are well approximated by a normal distribution whose mean and variance define a front wave that runs and widens exponentially fast in time outwards (see Appendix, section A2 for details):

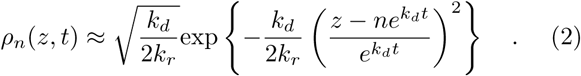

Equation (2) tracks the density of a single lineage through time and space. The whole system dynamics, however, is composed of different lineages competing to reach monoclonality. In addition, a given position can only contain a single cell, making a rigorous analytic treatment intractable. However, since the competition for a given position is neutral, one can make the approximation that the above derived densities reflect the likelihood that a given lineage occupies the considered position. In that context, the relative value between two different densities would therefore reflect the rate at which a given lineage would outperform the other. Consistently, if we consider all lineages competing, we conclude that the probability of lineage survival after a large time period can be derived directly from the normalization of the asymptotic densities, i.e.:

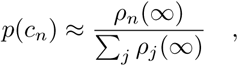

where, interestingly, *ρ*_*n*_(∞) ≡ lim_*t→∞*_ *ρ*_*n*_(*z, t*) is a constant independent of the position *z*. Importantly, this leads us to a well-defined, stationary probability of lineage survival:

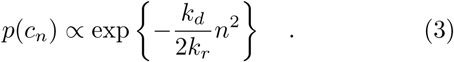

The above equation, which is a central result of the paper, predicts the probability that a cell starting at position *n* will “win the competition” and colonize the whole one-dimensional system (see Appendix, section A4 for details).

## II. FUNCTIONAL STEM CELL NUMBERS AND DYNAMICS IN THE STOCHASTIC CONVEYOR BELT

The prediction for the probability of long-term lineage survival is surprisingly simple, decaying as a Gaussian distribution as a function of position away the niche, with a length scale that is simply the amplitude of the stochastic fluctuations divided by the proliferation rate, 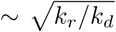 (see equation (3)). Intuitively, cells close to the origin have the highest chance to win and survive, whereas this probability drops abruptly for cells starting the competition further away, i.e. around *N*_*s*_ cell diameters away from the base, with:

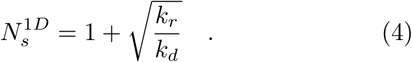

Note that the first term satisfies the boundary condition this at, in the case *k*_*r*_ = 0, the system has a single functional stem cell (located at the base) in 1D. Equation (4) thus implies that multiple rows of cells possess longterm self-renewal potential (as assessed for example, in a lineage tracing assay), emerging through their collective dynamics, and with a number that depends only on the ratio of the division to rearrangement rates (resp. *k*_*d*_ and *k*_*r*_). Although equation (4) is a 1D criterion, we show (Appendix Text, section B) that it holds and can be generalized in more complex geometries. In particular, in a cylindrical 2D geometry, we show the functional stem cell number would simply be the same number 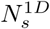 of cell rows (arising from the stochastic conveyor belt dynamics), multiplied by the number of cells per row (fixed by the geometry of the tissue). Therefore, this generalizes the results of Ref. [16], as we do not fix the stem cell number *N*_*s*_ explicitly, which rather emerges from an interplay between geometry and stochastic conveyor belt dynamics, together with the competitive dynamics being qualitatively different in the flow direction (Appendix Text, section A).

In spite of the approximations outlined above, stochastic numerical simulations of the model system show excellent agreement with equation (3)(see Fig. (1d,e)). We also note that although we model cells with identical intrinsic properties, the model can be extended be more complex cases, for instance with non-neutral cells (see below) or cells away from the niche going through irreversible differentiation. Moreover, although we have assumed here that positional rearrangements occur between two cells, more complex sources of noise can be considered, and lead to the same qualitative results. These include, for instance, post-mitotic dispersal, as seen during the branching morphogenesis of the kidney uteric bud [25] and where daughter cells can travel long distances outside the epithelium post-division, or correlated “tectonic” movements of the epithelium, where cells could collectively reposition relative to the niche, as proposed during mammary or gut morphogenesis [9, 21] (see Appendix section C for details).

We now turn to experimental data to test whether the proposed dynamics can help predict the number of functional stem cells in several organs, as well as the evolution of the survival probability with starting position of a clone. Although the division rate *k*_*d*_ is well-known in most systems considered, the stochastic movement rate *k*_*r*_ is harder to estimate, and can potentially vary widely, from rather small in intestinal crypts [13], to large in mammary and kidney tips, with extensive clonal fragmentation and random cell movements [9, 24].

### A. Predictions on clonal dynamics and survival

Intra-vital live-imaging provides an ideal platform to test the model, as it provides both knowledge of the starting position of a given cell as well as its clonal time evolution (whereas classical lineage tracing relies on clonal ensembles obtained from fixed samples). In small intestinal crypts, different Lgr5+ cells have been predicted to have very different lineage survival potential on the short-term, depending on their position within the stem cell niche, resulting in an effective number of stem cells smaller than the number of Lgr5+ cells [13, 30]. We thus reanalyzed quantitatively this dataset by plotting the survival probability of a clone as a function of its starting position *n* (see Fig. (2) after a given time period, *p*(*c*_*n*_, *τ*). We then compared this to a 2D stochastic simulation of the model (see Appendix section D for details). Importantly, we found a good qualitative and quantitative agreement between model and data, with the survival probability decaying smoothly with the starting position (see Fig. (2b)). The only parameter here was *k*_*r*_*/k*_*d*_ = 1, which fits well with short-term live imaging experiments and the idea of cell division promoting cell rearrangement [13].

**FIG. 2:**
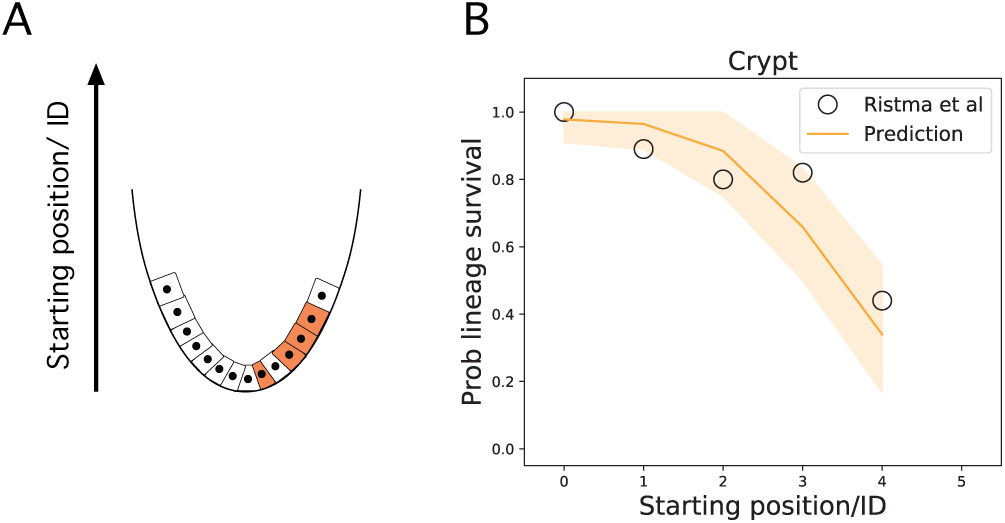
A/ Schema of the self-renewal of the crypt epithelia, showing the origin of the coordinate system at the bottom of the system. B/ The probability that a given cell remains within the system as a function of the starting position after a time lapse against the predictions of the conveyor belt dynamics for the crypt. Data corresponding to the probability that a lineage remains in the system for small intestinal crypt, reported in [13] depending on its starting position. The orange line represents the prediction of the stochastic conveyor belt dynamics, fitting well the data for *k*_*r*_*/k*_*d*_ ≈ 1. Shaded areas represent the confidence interval (1S.D.) of the prediction.

To back these simulations with an analytical prediction on stem cell numbers, the details of tissue geometry must be taken into account (with the number of cells per row *i* needing to be estimated, while the number of rows participating in the competition arising as an emergent property from the 1D model). A good approximation is based on that fact that the crypt can be abstracted as a hemispherical monolayer with radius *R* (measured in units of cell diameter) coupled to a cylindrical region (see Appendix, section B for details), so that one can get the number of stem cells, 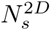, as:

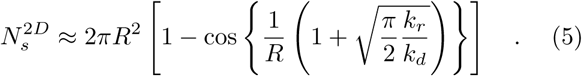

The above prediction is similar to the 1D prediction in equation (4), although it takes into account the effect of tissue geometry. With *k*_*r*_*/k*_*d*_ ≈ 1 as above, and estimating *R* ≈ 2 for the radius, our simple theory then predicts that the number of functional stem cells should be *N* ^2*D*^ ≈ 8, which agrees well with measurements of [13], as well as inferred numbers from continuous clonal labelling experiments [30], which is expected, as our model reduces to the 1D ring model of Ref. [16] for low *k*_*r*_*/k*_*d*_. We then sought to test the model further using a published dataset on embryonic kidney branching in explants [24], which has been recently noted to be a highly stochastic process, with neighbouring cells at the start of the tracing ending up either surviving long-term in tips or being expelled to ducts. Moreover, Ref. [24] observed extensive random cell intercalations, in addition to the previously described mitotic dispersal [25], where cells extrude from the epithelium post-division and reinsert at a distance of *d*_*c*_ cell diameters away. Importantly, these processes can still be captured as an effective diffusion coefficient *k*_*r*_ in our framework (see Appendix, section D for details). Specifically, knowing that the fluctuations may occur at each duplication, and that they imply a displacement up to *d*_*c*_ ≈ 2 − 4 cell lengths, we can estimate that 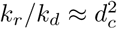 at the minimum (i.e. discounting other fluctuations). Note that the conveyor belt dynamics applies exactly as in the crypt: The only difference is that the reference frame from which the dynamics is observed now changes and, instead of taking the bottom of the organ as the rest reference, we take the newly created ductal cells produced by the tips. In that reference frame, the dynamics predicts a continuous elongation of the tips of the branches, as observed in reality (see Fig. (3a) and Appendix, section A for details).

**FIG. 3:**
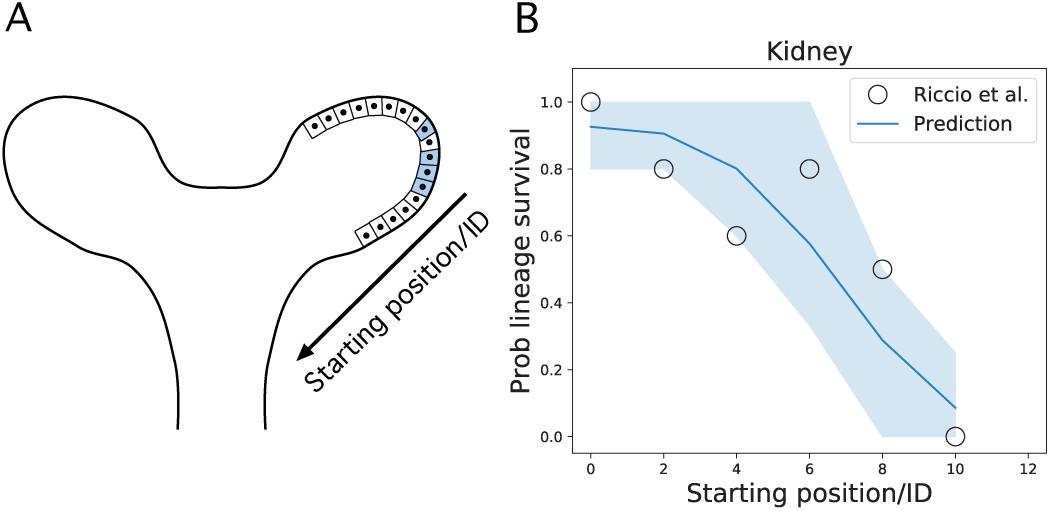
A/ Schema of the kidney tip during development. The conveyor-belt dynamics holds, the only difference is the reference frame: Whereas in the stem cell replacement model of the intestinal crypt the reference frame is the bottom of the gland, in the kidney and mammary gland, the reference frame is taken from the newly created ducts. B/ The probability that a given clonal remains within the system as a function of the starting position of the mother cell after a given time against the predictions of the conveyor belt model dynamics. Black circles represent real data points, obtained by counting the amount of cells of a given lineage remaining in the system (from Ref. [24]). We observe that the distribution is much broader, fitting well to the theory for a ratio *k*_*r*_*/k*_*d*_ ≈ 16 in kidney, much bigger that in intestinal crypt. Shaded area represents the confidence interval (1S.D.) of the prediction.

The above observation argues again that noise will play a key role in kidney tip cell dynamics. Strikingly, extracting from Ref. [24] the probability of survival as a function of distance from the edge of a tip, we found that the 2D simulations of our model provided again an excellent prediction for the full probability distribution (see Fig. (3b)), with cells much further away (compared to the intestinal crypt) having a non-negligible probability to go back and contribute. Again, the only fit parameter was the ratio *k*_*r*_*/k*_*d*_ = 16, which agrees well with our estimate of the noise arising from mitotic dispersal. Taking into account the full 2D geometry as above, and estimating in this case a tip radius of *R* = 3 − 5 cells, this predicts *N*_*s*_ ≈ 90 ± 10, which could be tested in clonal lineage tracing experiments.

These two examples show that the same model and master curve for the survival probability of clones can be used in different organs to understand their stem cell dynamics, and shows that ratios of relocation to advection *k*_*r*_*/k*_*d*_ can be widely different even in systems with similar division rates *k*_*d*_.

### B. Number of functional stem cells in the developing mammary gland

Next, we sought to test the model in the setting of mammary gland morphogenesis, where extensive cell movements have been reported within tips via intravital live-imaging [9], with rapid rearrangements occurring on time scales of a few hours (see Fig. 4 and Fig. S7 of the SI). In this case, however, tips cannot be followed for long-enough for survival probabilities to be inferred as in Figs. (2) and (3) for intestine and kidney, respectively. However, extensive clonal dispersion has been observed in quantitative clonal lineage tracing experiments during pubertal growth [9, 31], and we therefore sought to develop a statistical method to infer the value of noise from these experiments.

**FIG. 4:**
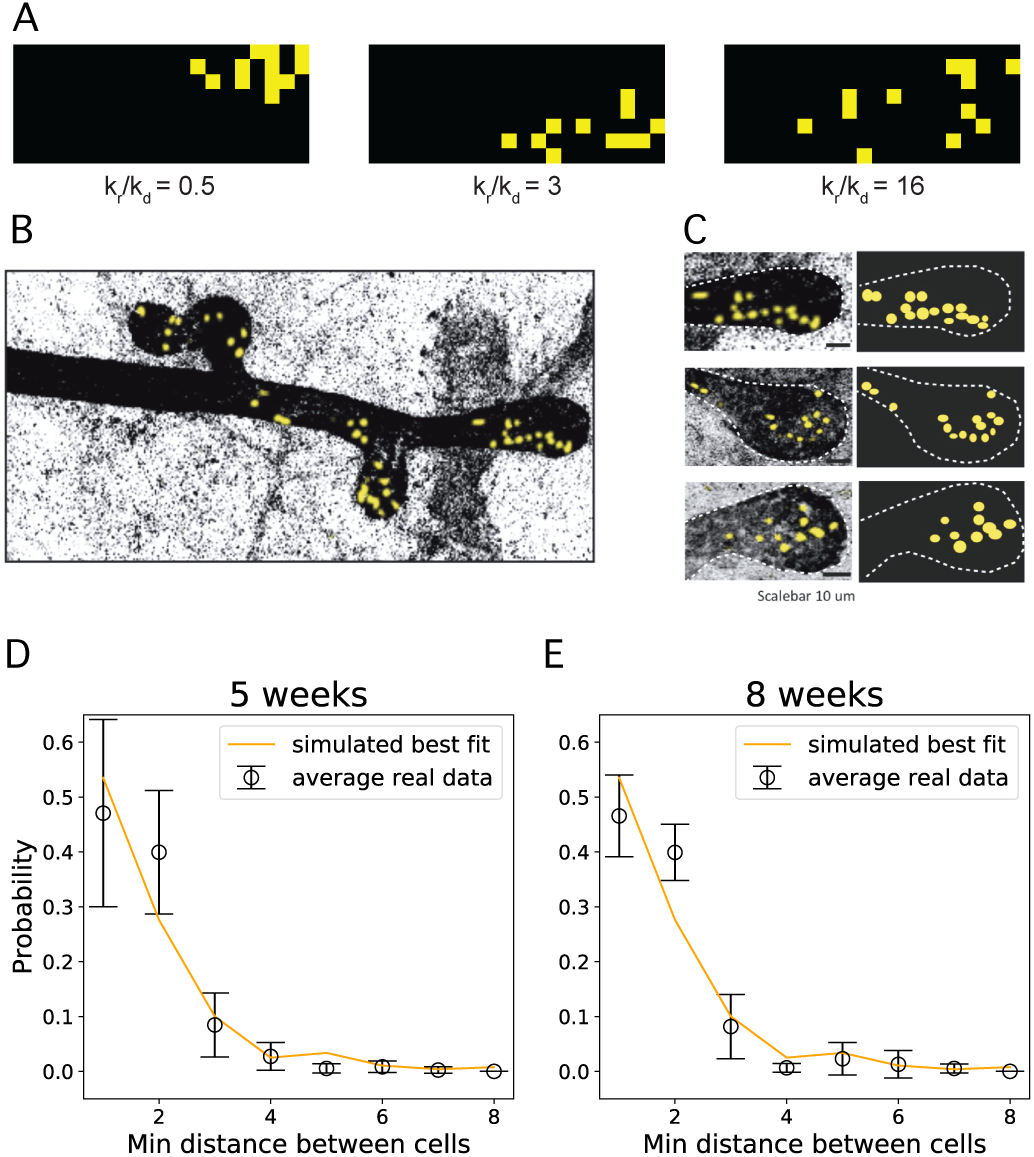
A/ Inferring the relation *k*_*r*_*/k*_*d*_ from the clone dispersion using a simulation of the stochastic conveyor belt dynamics in 2D. The distribution of distances of the closest neighbours is highly sensitive to the relation *k*_*r*_*/k*_*d*_. Here we show numerical simulations of fragmentation under constant *k*_*r*_*/k*_*d*_. B/ The tip of a developing mammary gland, where cells of a given lineage are coloured differently (yellow). Some dispersion of the clone due to random cell rearrangements is observed, as the clone is not perfectly cohesive. C/ Raw detail of the end buds of the tips forming the mammary gland (left) and reconstruction (right), where the lack of cohesiveness of the clone is clearly appreciated. The geometry of the end buds can be approximated by a hemispherical structure connected to a cylindrical one whose radius can be inferred to be around 2 − 5 cell diameters. D/ Closest distance dispersion curve obtained after 5 weeks. The curve has been obtained by running 2D simulations with different ratios and then fitting the closest distance distributions with the real data, obtained from analyzing different tips. Basal and luminal cells have been treated together for this analysis, as they do not show different behaviour at the level of the dynamics. E/ The same analysis for the closest distances distribution after 8 weeks. Both 5 weeks and 8 weeks showed a quite robust fitting at *k*_*r*_*/k*_*d*_ ≈ 3.

Turning back to published lineage-tracing datasets, where single mammary stem cells are labelled at the beginning of puberty (3 weeks of age) and traced until either 5 weeks or 8 weeks of age, clones in tips displayed extensive fragmentation, which is expected to be directly related to the ratio *k*_*r*_*/k*_*d*_ (see Fig. (4c,d)). We thus ran as above two-dimensional simulations of our stochastic conveyor belt model (see Appendix, section D, for details), using measured values of the tip width and length to set the geometry. As a metric for clonal dispersion, we then computationally measured for each labelled cell the distance to its closest clonal neighbour: for a fully cohesive clone, all cells should be touching and the distance to the closest neighbour should be always one cell diameter. Increasing the value of *k*_*r*_*/k*_*d*_ robustly increased the closest neighbour distance. We then performed the same measurements in the experimental data set, both for the 5 weeks and 8 weeks time points (see Fig. (4c,d)), but also for luminal and basal cell types separately, given the demonstrated unipotency of these cell populations in pubertal development [9, 31]. We found highly consistent results in all four cases (average closest distance of around 1.85 cell diameter) which allowed us to infer a ratio of (see Appendix, section D):

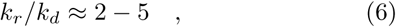

in mammary gland, emphasizing the importance of considering stochasticity in the conveyor belt picture. Indeed, we found that, with this fitting parameter, the model reproduced well the probability distribution of closest distances, both at the 5 weeks and 8 weeks time points (see Fig. (4c,d)).

In addition to this value, we must again pay attention to the geometry of the mammary tip, with basal cells forming a 2D monolayer (similar to the previous cases) while luminal cells form multiple layers in 3D within the tip. Assuming that the intercalation between cells occurs mainly at the same layer, the system of luminal cells in the tip of the mammary gland can be abstracted as *R* − 1 successive hemispherical 2D layers. Let us emphasize the dependence of N ^2D^, as defined by equation (5), on *R*, writing 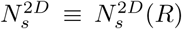. In that case, the amount of luminal stem cells can be inferred as:

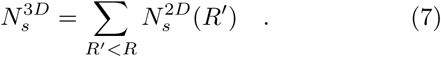

Taking the fitted range of *k*_*r*_*/k*_*d*_ ∈ (2, 5), together with an estimation of the radius of *R* = 5 ± 2, equation 7 then predicts that a number of luminal stem cell per tip of 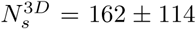, in good quantitative agreement with experimental estimates from lineage tracing of 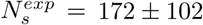 (mean ± s.d.) [9].

For basal cells, using the same parameters for a 2D monolayer, equation 5 predicts that 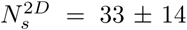, against empirical observations reporting an amount of basal stem cells of at least 15 [31], and 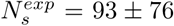 (mean ± s.d.) [9]. Although the prediction thus falls in the correct range, the under-estimation of basal stem cell number may be due to the highly anisotropic geometry of basal stem cells.

## III. DISCUSSION

The main objective of this study is to provide new insights to the question of whether stem cell function is a cell-intrinsic, inherited, property, or rather an extrinsic, context-dependent notion emerging from the collective dynamics of a tissue [2, 3, 7, 8]. To that end, we took a complementary standpoint to the one based on the classification of molecular markers and their potential functional role, adopting a purely dynamical/geometrical approach. In that sense, we tested to what extent the observed phenomena can be attributed to extrinsic, context dependent properties arising from the dynamics and geometry of the system. We analyzed in detail the dynamical properties of the process of tissue growth and self-renewal, taking into account the presence of stochastic cell rearrangements, cell duplication rates and tissue geometry. The combination of these ingredients gives rise to a complex reaction-diffusion process that can be abstracted as a stochastic-conveyor belt; that is, cells follow an exponential growth and are dragged along the tissue by an Ornstein-Uhlenbeck process with positive drift. Within this framework, we can predict several critical quantities, which emerge solely from the dynamics. Among others, the hierarchy of lineage survival, which has been shown to follow a Gaussian distribution that matches previously reported data, from kidney development and intestinal crypt renewal. From it, one can infer the effective size of the stem cell niche or functional number of stem cells, showing very good agreement with recent quantitative observations both in the crypt and in the development of the kidney and mammary gland. In the latter, we identified the traces of random rearrangements from the fragmentation of clones in lineage tracing data, obtaining an interesting steady ratio around *k*_*r*_*/k*_*d*_ ≈ 3, which is enough to allow realistic predictions of functional stem cell numbers in each mammary tip. Furthermore, the theory also makes non-trivial predictions on the survival probability of cells as a function of distance from niches, which we could validate both in intestinal crypt homeostasis and kidney morphogenesis, although the ratio of noise to advection *k*_*r*_*/k*_*d*_ differed by more than an order of magnitude in both systems [13, 24]. We finally evaluated the impact of the presence of mutant, fast replicating cells on the lineage survival dynamics, which were revealed to be strongly dependent on the number of functional stem cells in wild-type. This provides simple predictions that could guide further experimental work. It is worth emphasizing that these predictions rely only on the geometry of the tissue, and the ratio between the stochastic rearrangements and cell duplication rates.

We also emphasize that, although we have sketched here the simplest source of noise in cellular movements (random exchange of position in cell neighbors), our approach and results are in fact highly robust to different types of microscopic assumptions, and should thus be seen as representative of a general class of models for stem cell dynamics with advection and noise, rather than a specific microscopic mechanism. In mammary gland and kidney morphogenesis, direct cell-cell rearrangements are observed [9, 24], while kidney also displays mitotic dispersal [25], where noise arises from the randomness of cell re-insertion in the layer after division. Furthermore, on short-time scales, directed cellular movements have been observed in kidney tip morphogenesis, with Ret and Etv4 mutant clones being statistically overtaken by wild-type cells, leading to the proposal that Ret/Etv4 were involved in directional movement towards tips [24]. However, tips maintain heterogeneity in Ret expression through branching, arguing that cells must shuttle between high-Ret and low-Ret states [24]. In our model, short-term directional movements based on Ret signalling, which is itself variable would still result in long-term diffusion motion, effectively give an additional contribution to *k*_*r*_ on long time scales. Finally, we also envisioned a source of noise *k*_*r*_ which does not result in clonal dispersion. We show indeed that “tectonic” and collective movements of the monolayer away from the niche (see Fig. S5 of the SI), would again give rise to the exact same results on the shape of the survival probability as described here (again resulting in effectively higher *k*_*r*_). Such tectonic movements are particularly relevant in developmental settings, such as gut morphogenesis, where the shape of the intestine changes, for instance via villi bending, which displaces collectively cells from villi regions to niche/crypt regions [21]. This is also relevant for branching morphogenesis, as proposed in Ref. [9], because branching collectively repositions cells into tip regions due to global shape changes of the epithelium, and could be an additional source of noise explaining the high value of *k*_*r*_ we inferred in kidney morphogenesis, for instance.

What are the consequences of this intrinsic stochasticity, beyond the emergence of a structured stem cell region? We made the observation that the size of the stem cell niche, understood as the fraction of cell-lineages with non-negligible probability of colonizing the system, may differ from the dividing region. This result is relevant because it complements and enriches the concept of a stem cell niche. Moreover, it is worth pointing out that a positionally-defined stem cell potential, jointly with stochastic fluctuations, can serve as a protection mechanism against death or malfunctioning of the tissue. Indeed, the existence of fluctuations can give rise to the possibility of spontaneous replacement ensuring the healthy functioning of the tissue. The exploration of the consequences of such stochasticity and trade-offs is an active field of research in stem cell modelling, both in the context of cancer [35–39], but also for instance in ageing [40].

The proposed framework is in fact completely general and can be, in principle, applied to any tissue dynamics in which the replication occurs from a given spatial point, such that the growth can be projected in a given, well defined single coordinate. Nevertheless, this strong geometrical constraint is not always playing a predominant role, for instance in the context of an “open niche” such as spermatogenesis [41] or skin homeostasis [11] where most cells attached to the basement membrane form a 2D layer of equipotent progenitors. In spermatogenesis, competition instead occurs at the level of diffusible mitogen/fate determinants consumption [12, 41], which shares some conceptual similarities with our model. Outside of epithelial tissues, macrophages and fibroblasts have been shown to form a stable circuit of interactions, giving robustness to the density regulation of each population [42]. Therefore, our approach must be taken as part of a more general enterprise, namely, the role of the complex, global dynamics of the system in defining the functional stem cells. Indeed, understanding the interaction between the dynamics we propose here and the role of the molecular markers/functional differentiation dynamics *in vivo* would be a natural next step. For example, the prediction of the number of stem cells in the crypt, even though accurate with respect to live-imaging functional data, does not explain why Lgr5 levels display a clear on/off pattern inside vs. outside the crypt, and reversal to a stem cell fate is not always a fast process dependent solely on niche cues. Therefore, a natural follow up should explore the relation between the emergent properties of the dynamics and the molecular markers, jointly to their functional differentiation [43, 44]. In addition, more complex geometries, and with that, other tissues, could be explored within our framework.

## IV. ACKNOWLEDGEMENTS

We thank all members of the Hannezo, Simons and Van Rheenen groups for stimulating discussions.

## Appendix A: Basic dynamics

The simplest abstraction of our system is composed by a column of cells arranged in a finite segment [0, *N*], in which the length unit is scaled to the average length of a cell. Each cell divides at constant rate *k*_*d*_. We consider that the system can only grow towards the positive axis and that the density of cells remains constant. This division creates a positive push up force because newly born cells occupy upper adjacent positions in the array, pushing the cells that were there to higher positions. As long as a cell reaches the position *N* and is pushed, it disappears from the system. Both intestinal crypts and elongating mammary or kidney tips can be described by this dynamics, up to a change of referential frame: intestinal crypts expel differentiated cells with the crypt base staying stationary, while differentiated cells stay in place while the tip moves forward in the case of mammary gland or kidney development. The coordinate *x* or *z* through which the dynamics take place will depict only the distance to the origin of the organ, without assumptions on the global movement of it: In the crypt that will represent the distance to the bottom, in the case of the kidney or the mammary gland, the distance to the tip –see figure (5). In addition, the position of the cells can fluctuate stochastically at rate *k*_*r*_ (either via local cell-cell rearrangements, or more global movements of cells relative to the niche, see sections below for more details). Importantly, the results that we derive, as described below, are highly generic to different types of assumptions on the microscopic dynamics, as long as the basis features of advection and rearrangements exist in a system (sketched in Fig. 1 of the main text).

**FIG. 5:**
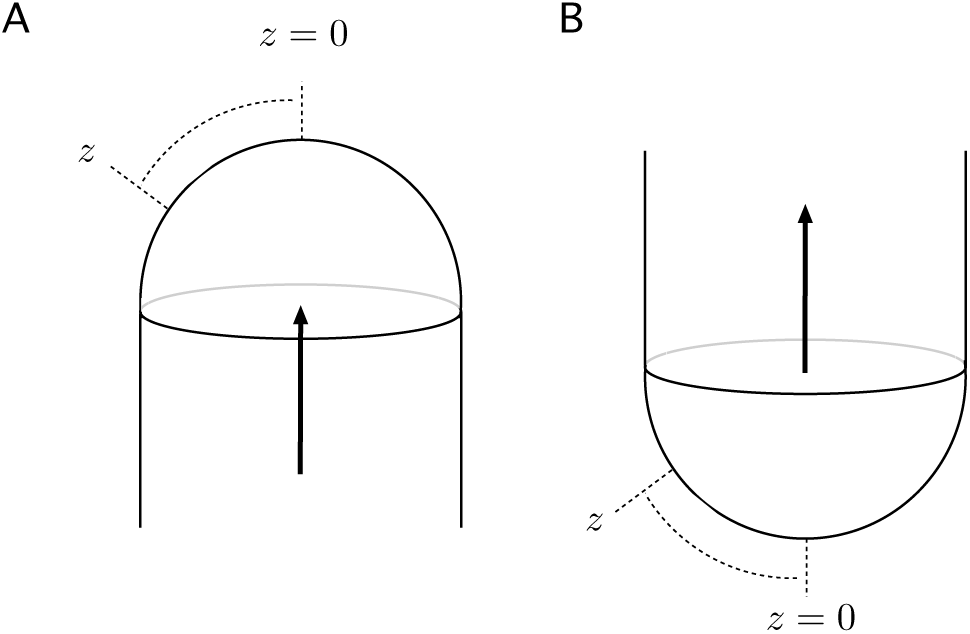
Equation (A1) is defined in terms of the coordinate *z*, that is, the distance in unit length of cell lengths from the origin of the organ, and has no assumptions on the global movement of it. A/ In the case of both the mammary gland and kidney development, the organ generates new cells that remain at rest, whereas the tip of the organ moves on. B/ In the case of the self-renewal of the crypt, the bottom of the system is at rest and cells are running upwards, whereas the organ does not move. In consequence, equation (A1) can describe both the dynamics of intestine crypt self-renewal and the mammary gland and kidney development.

Before proceeding with the details of the derivations and numerical computations, we present an overview of the strategy followed to derive the result. The aim is to convey a bird-eye view over the whole strategy, and to highlight the assumptions underlying the approach. The modelling and conceptual steps are the following:

1. The first step is to define the conveyor-belt dynamics at the single cell level. The probability that a cell that started at position *k* is at a given position of the organ at time *t*. This dynamics obey a Ohrnstein-Uhlenbeck process with positive drift. The evolution of the probabilities is governed by the following Fokker-Planck equation:

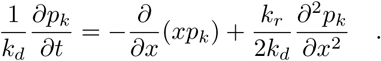
2. We then add the *reaction* term, that is, the cell, aside the random fluctuations provided above, also divides at a certain rate. The reaction-diffusion equation governing the density of lineage *c*_*k*_ is:

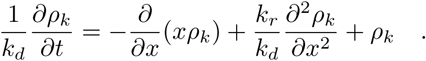
3. In spite the above system has no stationary solutions, the density of the lineage *c*_*k*_ at a given position of the organ goes to a constant over the whole organ as *t* → ∞. We call this density *ρ*_*k*_(∞).
4. We *assume* that *ρ*_*k*_(∞) is proportional to the probability that the lineage *c*_*k*_ will colonize the whole organ. Therefore:

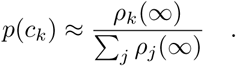
5. This probability turns out to be a gaussian-like distribution depending only on the ratio *k*_*r*_*/k*_*d*_:

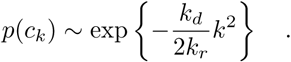
6. Once the 1D problem is solved, we go to more general geometries. As long as the push up force is projected towards a single coordinate of the system, the general reaction-diffusion reads:

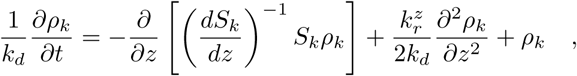

where:

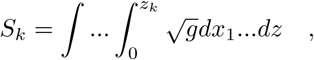

*g* is the determinant of the metric tensor, *z* is the coordinate over which the push-up force due to the fluctuations is projected and 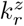 the explicit value of these fluctuations projected over the relevant coordinate *z*.
7. After some approximations, we reach a differential equation that can be integrated, for the case of geometries corresponding to cylinders coupled to hemispheric regions, as is the case of the organs studied:

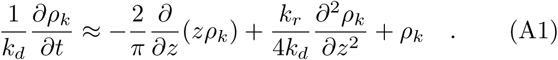
8. We finally study the cases where different *k*_*r*_*/k*_*d*_ rates are present in the same system.

### 1. Single cell dynamics

This process is a mixture of Brownian motion with a given amplitude *k*_*r*_ and a drift parameter that depends on the position. In the continuous limit, the position of the cell, *x*, can be described as a random variable satisfying the following stochastic equation:

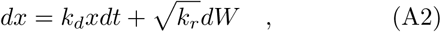

being *dW* the differential of the standard Wiener process with mean 0 and variance 1. In other words, we are describing a kind of *Ornstein-Uhlenbeck* process, with positive drift *k*_*d*_ [26, 27]. The above described stochastic process has no stationary solutions, which is in agreement with the conveyor belt-like dynamics of the systems under study: all cells will sooner or later be pushed out from the system (in the case of the crypt, or will be left behind, as in the case of the mammary gland development, note that for finite systems, this will consist of a single remaining winner lineage colonizing the whole tissue). One can still compute the time-dependent solutions, considering initial conditions *t*_0_ = 0, *x*_0_ = *δ*(*x* − *k*) and natural boundary conditions^1^ as follows: First, we observe that:

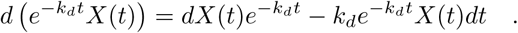

Then, multiplying both sides of equation (A2) by 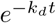, and after some algebra, one finds that:

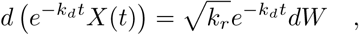

leading to:

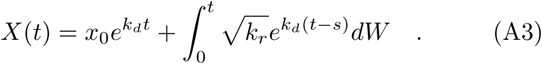

The integral is a standard stochastic integral with respect to a Wiener process. According to *Ito’s isometry* [45] one has that the law governing the random variable described by the integral is a normal distribution **N**(0, *σ*^2^(*t*)). In our case this reads:

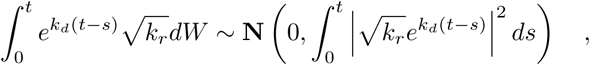

which means that he explicit form of *σ*^2^(*t*), is thus given by:

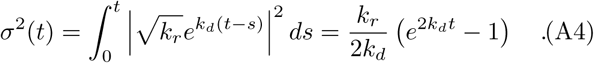

Finally, from equation (A3), we conclude that the time dependent mean, *µ*(*t*), is:

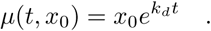

By setting

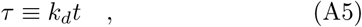

and *x*_0_ = *k* as the starting position, one gets the time evolution of the probability distribution of for the position of the random walker which started at *k*:

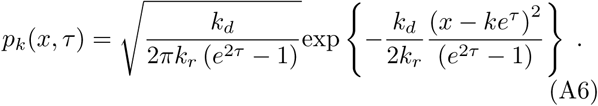

In words, the solution is given by random variable following a normal distribution whose mean and variance run exponentially fast in time through the positive axis. In Fig. (6a) of this Appendix, we plotted some snapshots of this time dependent probability.

**FIG. 6:**
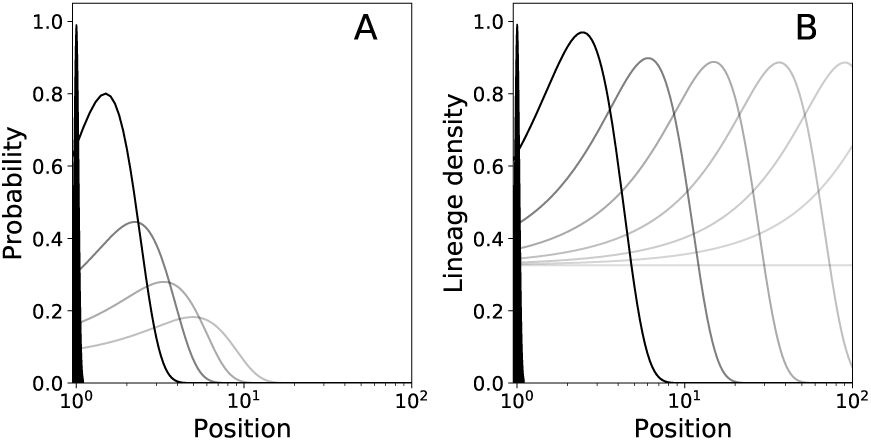
Temporal evolution of the stochastic conveyor belt dynamics (black to grey indicates time). A/ Evolution in time of the probability for cells starting at a given position to occupy the location *x* according to the theoretical prediction given by equation (A6). Observe that the dynamics does not run to a stationary state, so all cells will eventually abandon the system with probability 1 as long as time grows. B/ Evolution in time of the density of a lineage starting in the same position across all positions, according to the solution of the reaction diffusion equation (A8) given in equation (A11). Observe that the reaction diffusion dynamics displays a front that runs exponentially towards the outside of the system. However, we observe that the density reaches a non-zero stationary value, that is proportional to the probability of the lineage to remain and colonize the whole system. The initial black, elongated triangle at position 1 shows the initial conditions i.e., *k* = 1, and *k*_*r*_ = 1, *k*_*d*_ = 1. These values have been chosen only for the sake of clarity.

### 2. Lineage dynamics

The Fokker-Planck equation accounting for the probability that a given random walker starting at *k* will be at *x* at a given time *τ* –as defined in equation (A5)–, for the stochastic process described by equation (A2), *p*_*k*_(*x, τ*), is given by:

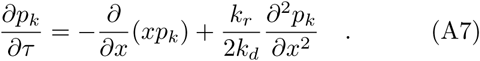

If we want to study the *density* of cells of the lineage started by cell *k* in a given position, *ρ*_*k*_(*x, τ*), we have to take into account that, in a given time step, *ρ*_*k*_(*x, τ*) new cells of the lineage will emerge in such a position. Therefore, to describe the whole process, we need a diffusive part, given by the differential operator 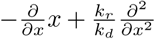 of equation (A7) and a reactive part, given by *ρ*_*k*_(*x, τ*). To derive the reactive part, we take into account the following reasoning: If the density of a given lineage at position *x* at time *τ* is *ρ*_*k*_(*x, τ*), one expects that, in average, new *ρ*_*k*_(*x, τ*) cells will emerge within this interval in a time unit –as it is standard in the exponential growth. So, one has that the equation accounting for the time evolution –in units of *τ* = *k*_*d*_*t*– of such density is:

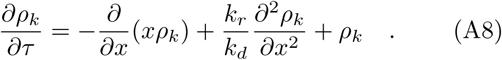

The above equation is a reaction diffusion equation whose diffusive part observes a Ornstein-Uhlenbeck processes with positive drift and whose reaction part is given by an standard exponential growth. Knowing that the solution of equation (A7) is given by *p*_*k*_(*x, τ*) as defined in equation (A6), the solution of equation (A8), with initial conditions *ρ*_*k*_(*x*, 0) = *δ*(*k* − *x*), is given by:

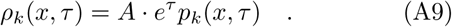

We observe that, according to equation (A9), the term *e*^*τ*^ *p*_*k*_(*x, τ*) can be safely approximated by

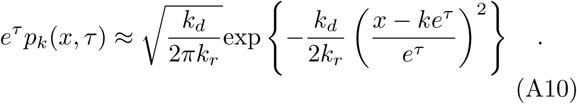

### 3. Determination of the integration constant

We observe that, even though random fluctuations in cell positions are allowed, epithelial tissues remain confluent [46], and maintain a constant density at homeostasis. This imposes that, in the limit of a continuous array of cells and for *τ* → ∞, the overall density of cells of the site *x* must stabilize to 1, and must not depend on the position *x*. We will call that condition the *confluent tissue condition*. Consistent with the above reasoning, we observe that:

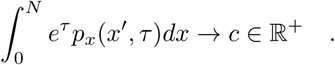

To determine the constant *A* of equation (A9), we impose the *confluent tissue condition*. According to the above equation, this is satisfied by:

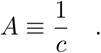

Let us remark that we consider *x* ∈ [0, *N*]. Then, knowing that, in this case:

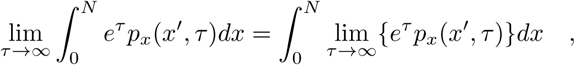

we compute the limit, taking into account equation (A10):

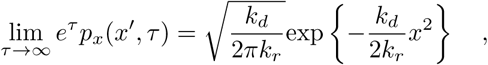

which is a gaussian with zero mean and *σ*^2^ = *k*_*r*_*/k*_*d*_. so that:

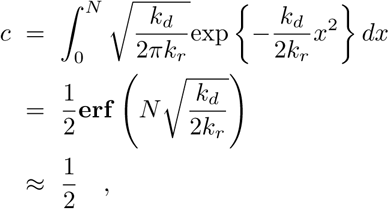

where **erf** (*x*) is the *error function*. The approximation *c* ≈ 1*/*2 holds as soon as *N* ≫ 1. Therefore, in that approximation, *A* = 2.

By setting the integration constant *A* = 2, the process fills up space in a correct manner that is: The density of cells is always conserved as 1 cell × length unit, as expected for a confluent homeostasic tissue. The final expression for *ρ*(*k, τ*) will thus be:

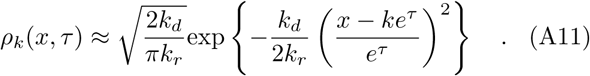

In Fig. (6b) of this Appendix we plotted some snapshots of this time dependent density.

### 4. Lineage survival probability

The first task is to demonstrate that the above dynamics leads, in the long term, to a system composed by descendants of the same cell, i.e., that only a single lineage is present in the system for *τ* ≫ 1. In the case of a 1D system defined over the interval [0, *N*] with *k*_*r*_ = 0, this result is trivial, since the lineage of the cell located at the bottom of the system will occupy the whole system whenever it has divided enough times to cover all the positions from 0 to *N*. In the case *k*_*r*_ > 0, the strategy to see that it runs towards monoclonality is the following: Let Ω_*N*_ be all the potential configurations the system can have in terms of lineage configuration whenever the system has 2, 3, …, *N* lineages alive. That is *σ* ∈ Ω_*N*_ will be a sequence of *N* numbers each labelling the lineage the cell in a given position belongs to:

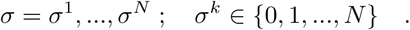

In units of *τ* and given a configuration *σ* of lineages, for 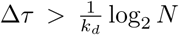 all the cells will potentially have produced more than *N* new cells. Therefore, the probability that a lineage has been expelled by the system will be larger than zero. Let us call this probability *p*(↓, Δ*τ*). Let us define 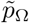 as:

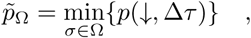

that is, the configuration for which the probability of losing a lineage after Δ*τ* steps is minimum. By construction, we know that from whatever configuration containing more than a single lineage has a non-zero probability of losing at least one of them after Δ*τ*, so:

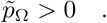

The probability of *not* losing any lineage whatever the configuration of the system after *n* steps of duration Δ*τ, p*(=, *n*Δ*τ*) will be bounded as:

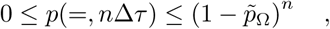

and the probability of losing *at least one* lineage will be bounded as well as:

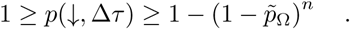

In consequence,

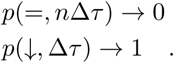

Since the loss of a lineage is a completely irreversible process, the above equation tells us that it is expected that the system will lose lineages until only one survives. Note that for a 2D geometry but *k*_*r*_ = 0, the system reduces to the stochastic voter model along a 1D ring (the *N*_0_ cells at position 0) which was proposed and tested experimentally in Ref. [16], and where the process of monoclonal conversion is diffusive, occurring on time scales of 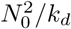.

Now that we know that the system will reach monoclonality, the next question is to ask which lineage will win the competition. In particular, we are interested in the probability that a given lineage colonizes the whole system as a function of the position of the cell that defined the lineage at *t* = 0. To that end, we first compute the asymptotic lineage density, *ρ*_*k*_(∞), that reads:

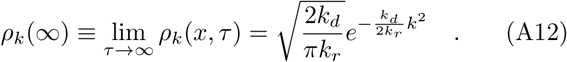

We observe that this density is independent of *x* –see Fig. (6b) of this SI. Now we make the following assumption: The competition between cells at different levels term is absorbed in a mean field approach by the drift push-up force. In this context, we conclude that the probability of lineage survival *p*(*c*_*k*_) –that is, the probability that the whole tissue from 0 to *N* will be occupied by cells of the lineage *k*– can be derived directly from the normalization of the asymptotic densities *ρ*_*k*_(∞):

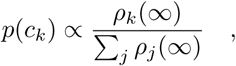

which leads to:

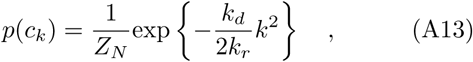

being *Z*_*N*_ the normalization constant, namely, 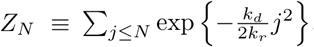.

## Appendix B: Dynamics in more general geometries

In general we will assume that there is a coordinate *z* over which the displacement induced by the division takes place. All the dynamics will be, in consequence, studied from its projection over this coordinate –see Fig. (7) of this Appendix for the special cases of hemispheric and spheric geometries. In the case of a 1D-system, as the one described above, this coordinate is the length, *x*. In the case of a hemisphere, assuming that the push-up force is exerted from the bottom pole, this coordinate is the arc length defined from the position of the cell to the bottom pole itself –see Fig. (7a) of this SI. To gain intuition, consider the surface of the hemisphere with radius *R*: The cells at the bottom pole divide and push the ones on top of them up through the surface. The cell under consideration is located at whatever position defining an arc from the bottom pole equal to *z* = *Rφ*_*k*_ = *z*_*k*_, where *φ*_*k*_ is the polar angle, meaning that there is an arc of *k* cells from the given cell to the pole of the hemisphere –see Fig. (7a) of this SI. The successive divisions of cells located at *z*_*i*_ < *z*_*k*_ will result into a net displacement along the angular coordinate *φ* of the cell located initially at *z*_*k*_ = *Rφ*_*k*_, going from 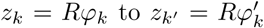, with 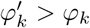. The linear displacement along the surface will be 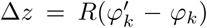. Displacements along the other coordinate will have no effect in the push up force.

**FIG. 7:**
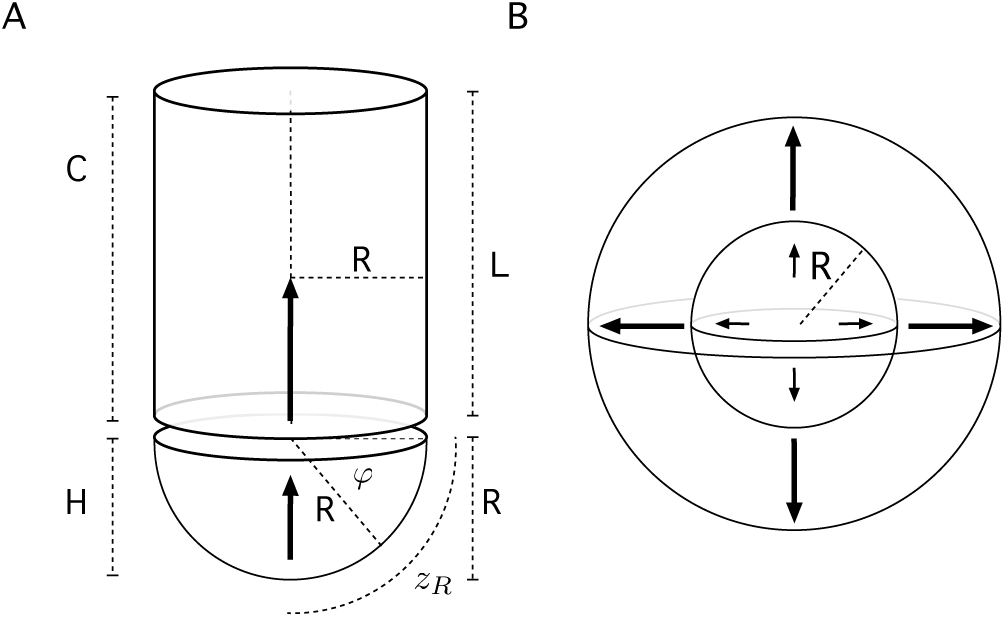
A/ Schematic characterization of the structure of the crypt as a hemispherical region *H* coupled to a cylinder region of the same radius, *R*. B/ The expansion of the tissue in a 3D abstract setting where there is radial symmetry. The growing of the inner cells creates a push up force. In addition, the stochastic fluctuations in the position determine the probability of lineage survival as a function of the starting point, as in the case of low dimensional approaches.

### 1. Drift term

The push-up force or drift term will be described by the function *h*(*z*), and will be defined as:

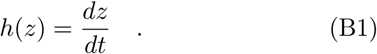

In the case of a 1D-system, as the one described by equation (A8), one has that *h*(*z*) = *k*_*d*_*z*. To properly study this dynamics over mor general geometries, let us consider a Riemannian manifold equipped with a metric tensor **g**, with components *g*_*ij*_ [47]. Crucial to our purposes is the property of *local flatness* [47, 48]. Roughly speaking, this implies that, for small enough regions of the manifold, the geometry has euclidean properties. Let us consider that the push up force due to duplications has an origin and is exerted along the direction of a single coordinate *z* as well. As we did above, the surface/volume units are given such that an average cell has a surface/volume of 1 in the corresponding units. Consider the starting position of our cell to be *z*_*k*_ along the coordinate *z* along which the displacement due to the push up force takes place. If the other coordinates are given by *x*_1_, …, *x*_*n−*1_, the surface/volume encapsulated below this position is given by:

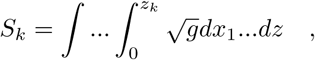

where *g* is the determiner of the metric tensor, i.e.:

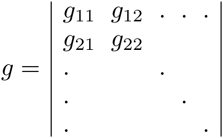

Cells are assumed to divide at rate *k*_*d*_. That implies that *k*_*d*_*S*_*k*_ new cells will be produced *below* the cell located at *z*_*k*_. This will create an extra surface/volume of:

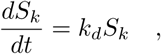

that will project into the coordinate *z*. Using that:

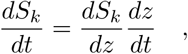

and, then, equation (B1), one can find the general expression for this projection, which reads:

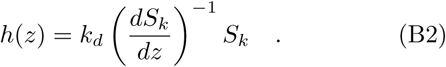

### 2. Projection of the fluctuations

We are only interested on the projection of the dynamics over the coordinate *z* along which the system grows, as in the other coordinates the competition is neutral and has no net effect in the lineage survival statistics. If the reported fluctuations are *k*_*r*_, we will refer to the projection to the coordinate *z* as 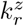. In general, 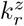 will be a function 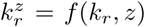. According to the above results, we will have that the general equation for the evolution of cell lineage densities along the coordinate *z* will read:

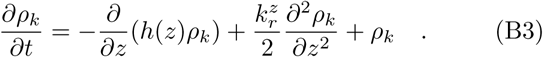

In the case *h*(*z*) can be approached as a linear function, i.e., *h*(*z*) ∼ *ak*_*d*_*z* and 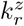 as 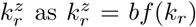, one can reproduce the reasoning provided to derive the lineage survival probability, equation (A13), and obtain:

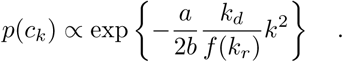

To gain intuition, imagine that we report experimental fluctuations of amplitude *k*_*r*_ –see Fig. (8) of this SI. That is, in a time unit, the cells move randomly over the manifold *k*_*r*_ steps. We are in a 2D isotropic, locally flat surface with generic orthogonal coordinates *y, z* –for example, *R*× the azimuthal angle *θ* and *R*× the polar angle *ϕ* over a sphere surface. The amplitude of the fluctuations after time *t* is known to be 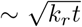, a distance defined over the surface. In the case we consider the projection over the coordinate *z*, thanks to the local flatness [48], assuming that 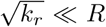 and using only symmetry reasonings, one has that since the displacement is given by 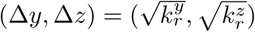:

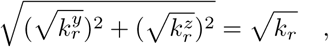

and the fluctuations are isotropic, then:

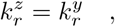

and the only solution to the above problem is that:

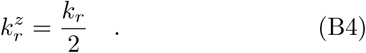

**FIG. 8:**
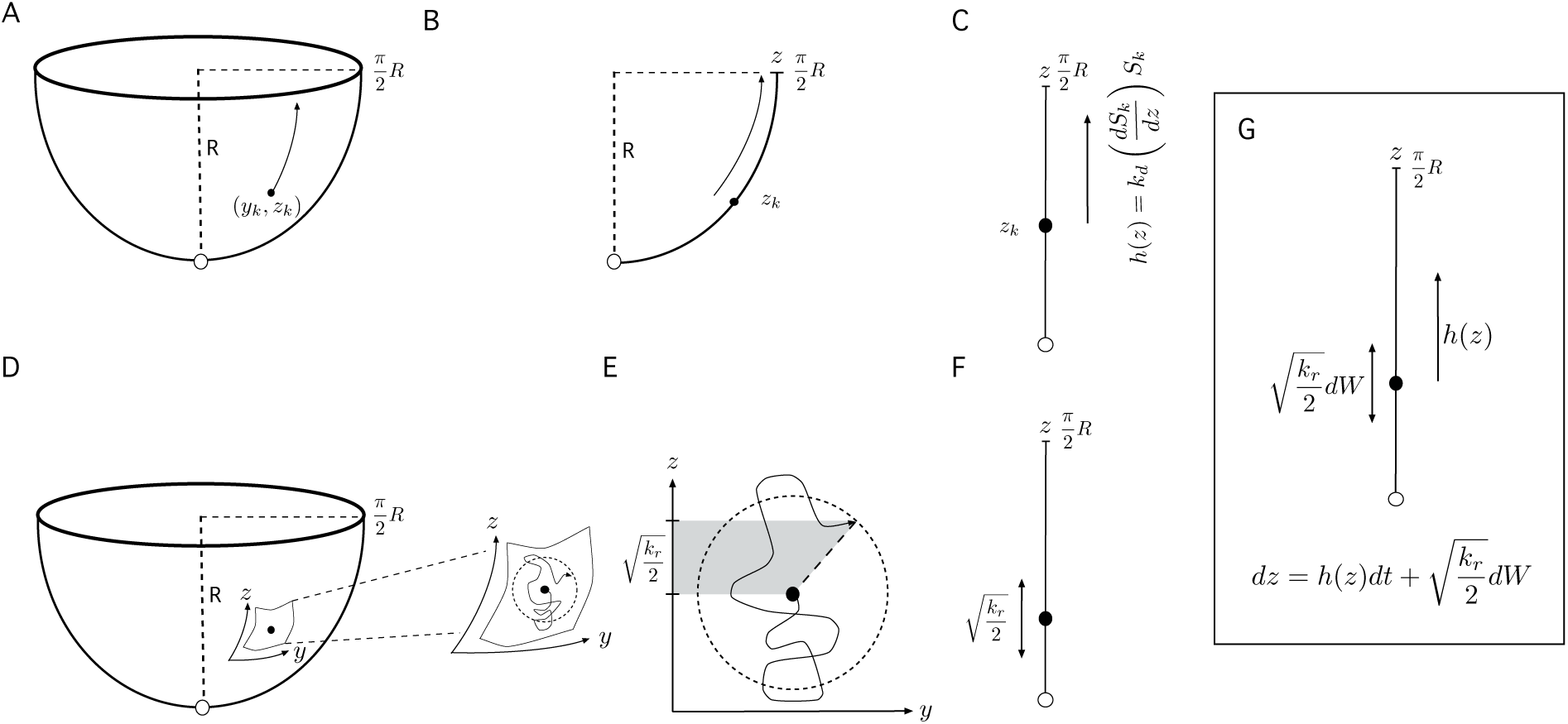
Constructing the dynamical equation for a general kind of manifold and projecting it onto the 1*D* coordinate system along which the push-up force is exerted. A/ A cell is located in a point in a manifold –in that case, a hemisphere–, described by two orthogonal coordinates (*y*_*k*_, *z*_*k*_). In this case *y*_*k*_ = *Rθ*_*k*_, where *θ* is the azimuthal angle, and *z*_*k*_ = *Rϕ*_*k*_, where *ϕ* is the polar angle. B/ the push up force due to duplication is only exerted along the *z* coordinate. C/ *h*(*z*) is the drift term that enters the equation, and refers to the amount of new surface that has been created below the point *z*_*k*_ in the *z* coordinate that results in pushing up the cells above. D/ the same point (*y*_*k*_, *z*_*k*_) also observes fluctuations due to random noise. In particular, the rate of these fluctuations is externally reported as *k*_*r*_. E/ Thanks to the local flatness property, if 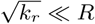, then the fluctuations take place locally in a flat space, and the average distance from the starting point, after a unit time interval, will be 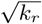. Since the coordinate system *y, z* is orthogonal and locally flat, and the random fluctuations occur isotropically in space, the projection of the fluctuations over the coordinate 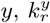, will be the same than in the coordinate 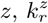. Since 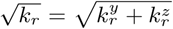, the only solution for this projection is that, 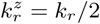, as described in F/. G/ Combining C/ and F/ we have that the growing process can be described along the *z* coordinate as a Ornstein-Ü hlenbeck process with push up force *h*(*z*) and random fluctuations 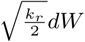, where *dW* is the standard Brownian motion with mean 0 and variance 1.

In the case we are dealing with a spherical surface, we are projecting the fluctuations over the polar angle *z* = *Rϕ* – see Fig. (8) of this SI. The stochastic differential equation that will describe the movement of a single cell in this manifold will be, for 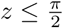:

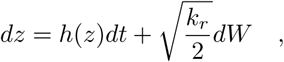

where *dW* is the differential of the standard Brownian motion with average 0 and variance 1 in one dimension.

### 3. Evolution of densities: Hemispherical approach

Let us now consider a detailed version of the geometry of the crypt. This consists in a half sphere, *H*, whose arc length from the bottom pole to the end is is 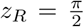 *R* coupled to a cylinder *C* of length *L* and radius *R*. The cells populate both the surface of the hemisphere and the cylinder. The push-up force is directed towards the top of the cylinder –see Fig. (7a) of this SI. In the arc that goes from the bottom pole to the end of the hemisphere there are *z*_*R*_ cells. Again, the units are given considering the average size of the cell as the length/surface/volume unit. Therefore, the cells will be labelled in terms of the geodesic distance over the hemisphere to the bottom pole. The density of the lineages will be given by:

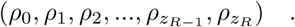

Since the coordinate *R* is constant, the only dynamically relevant information will come from the angle *ϕ*. Each position *k* in the arc (0, 1, 2, …, *z*_*k*_, …, *z*_*R*_ − 1, *z*_*R*_) describing the initial point of a cell lineage can be rewritten as:

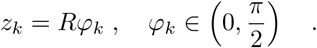

i.e., 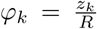. The metric tensor for this hemispheric surface is [48]:

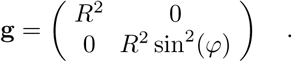

Computing the determiner of **g**, *g*:

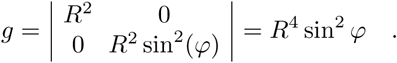

one can compute the surface element as [48]:

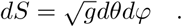

In consequence, the area under the position of the cell *k* in the in the hemisphere *H*, located at the arc position *z*_*k*_, will be:

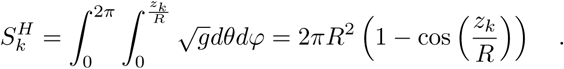

By direct application of equation (B2), we have that the push-up force inside the hemisphere *H* is given by:

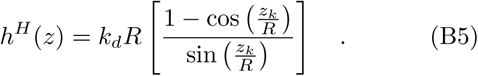

Finally, from equation (B4) we know that –see also Fig. (8) of this SI:

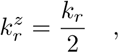

leading, according to equation (B3) to the general dynamical equation for 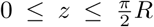 in a hemispherical surface to be:

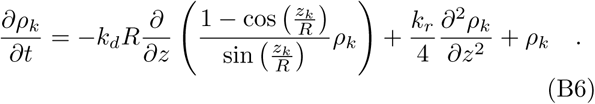

The above equation is difficult to deal with. However, we observe that in the region of interest, 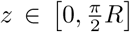, equation (B5) can be approximated as:

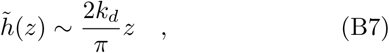

leading to an error bounded as:

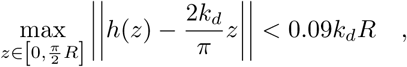

according to numerical tests. With this approximation, we have that equation (B6) can be rewritten approximately as:

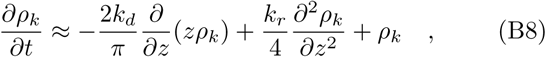

which is the general kind of Ornstein-Uhlenbeck equations we have been working so far.

### 4. Coupling to a cylinder

In the case the cell is at the position *k* in the cylindric region *C*, the area under it will be given by 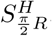, the area of the whole hemisphere, and the remaining surface due to the cell is the cylinder. Knowing that for the cylindric coordinates 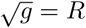, then:

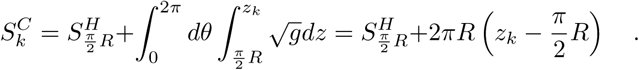

Completing the picture, the push force felt by a cell in the cylindric region *C* is given by:

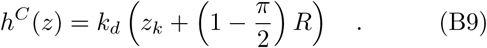

It is easy to check that:

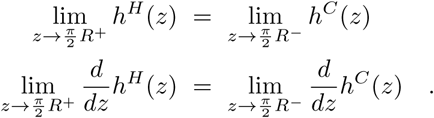

Therefore, one can define a function, *h*(*z*) as:

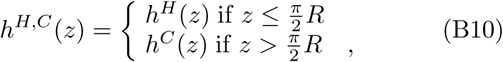

which is always well defined. In addition, the projection of the fluctuations will be the same in both regions, since the only relevant property is the local dimension, which in both cases is 2, leading to a 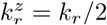. Consequently, equation (B3) can be rewritten consistently for all the hemisphere/cylinder system as:

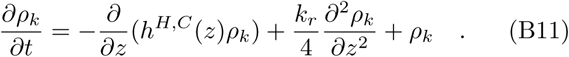

### 5. Lineage survival: Detailed geometry approach

To guess the probability of lineage survival, we restrict ourselves to the hemispheric region *H* of the crypt. As discussed in section B of this Appendix, the existence of linear functions approximating the drift and fluctuation parameters of the general reaction-diffusion equation (B3) leads to gaussian-like lineage survival probabilities. According to the approximations leading to equation (B8), we have that, considering the hemispheric region of the crypt:

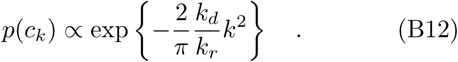

### 6. Number of stem cells in the system

To finish with, we compute the size of the emerging stem-cell region. The fluctuations will project over the surface from the bottom pole to the cell located at:

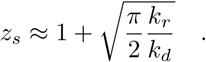

The position of this cell will define an angle of

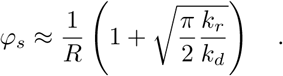

Therefore, the amount of cells below this angle, the stem cell number will be:

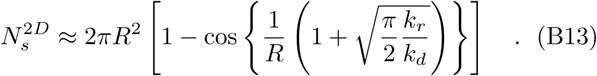

### 7. Higher dimensionality with radial symmetry

Imagine that our tissue can be abstracted as something isomporphic to a disk or sphere. In consequence, we will only consider the *r* coordinate –see Fig. (7b) of this SI. Let the units of volume be in number of cells. That is, we assume that, on average, the volume of cells is around 1. The fluctuations can occur in all directions with equal probability. Therefore, if we know that the rate at which fluctuations occur is *k*_*r*_, following the same reasoning we used in section B, we can approximate the effective rate projected towards the *r* axis by:

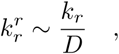

where *D* is the dimension of the volume. The diffusion term in the reaction diffusion equation will be given by:

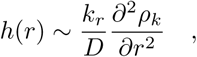

Now we compute the drift term. For that, we apply directly equation (B2). In the case of a 3D sphere growing from the centre, we have, if *V* (*r*) is the volume encapsulated by the surface of radius *r*, the displacement of a cell at this position will be determined by:

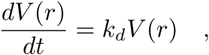

If 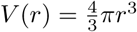, then, according to the above equation:

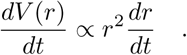

Thus, the push up force will, accordingly, result in a radial displacement of:

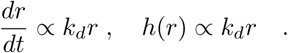

The above speed will define the drift term *h*(*r*). Again, this will lead to a gaussian distribution of lineage cell survival and, therefore, to a well defined stem-cell region.

## Appendix C: Noise in the stochastic conveyor-belt from “tectonic” epithelial movements

In this section, we explore an alternative source for noise in determining the number of functional stem cells, i.e. the possibility of global rearrangements of the epithelium relative to the optimal position (bottom of the crypt/edge of the tip –see Fig. (9a) of this Appendix for a sketch. This is motivated by experiments in intestinal morphogenesis or mammary morphogenesis, where global three-dimensional bending of the epithelial modifies the location of the niche (see main text).

**FIG. 9:**
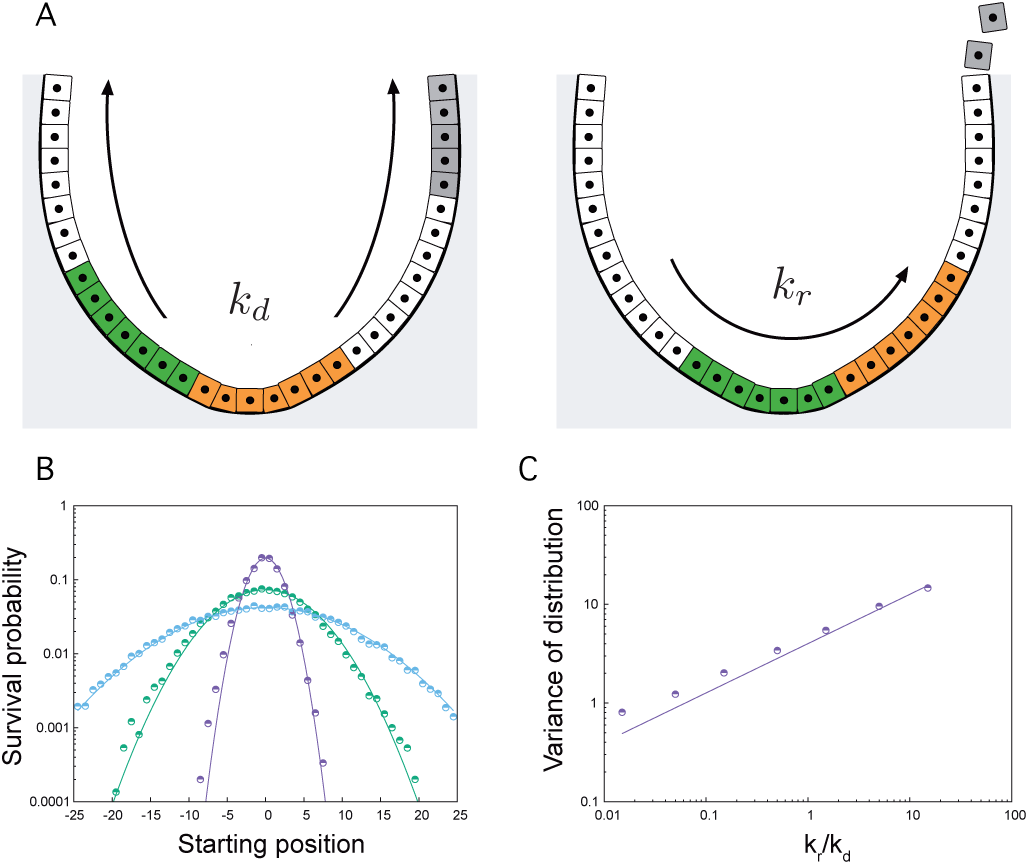
A/ Schematic of the model of stochastic conveyor belt with tectonic movements: cell division can occur for every cell, which pushes all cells above, but repositioning relative to the bottom of the crypt/tip can only occur via global movements of the layer, at rate *k*_*r*_ (two clones shown competing before and after a movement). B/ Computational predictions of the 1D stochastic conveyor belt dynamics in the presence of tectonic movements, with increasing rates *k*_*r*_ (purple to blue), in terms of survival probability as a function of starting position of the clone. All curves are very well fitted by normal distributions, as expected by our model. C/ Variance of survival probability as a function of starting position (i.e. functional stem cell number) as a function of the tectonic movements rates normalized by division rate *k*_*r*_*/k*_*d*_ (dots), and theoretical prediction (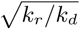, continuous line) from the stochastic conveyor belt model, showing that the system undergoes the same dynamics as for random stochastic intercalations.

Importantly, performing full stochastic simulations of this process in one-dimensions revealed a strikingly similar paradigm compared to the version of the model introduced in the main text, with survival probabilities decaying as normal distributions away from the central, optimal position for survival – see Fig. (7b) of this SI. Moreover, the variance of these probabilities, which define the number of functional stem cells, also scale as *k*_*r*_*/k*_*d*_ as expected in the model from the main text – Fig. (7c) of this SI.

This confirms that such tectonic movements can also be described in our coarse-grained model, simply renormalizing in long-term dynamics the intensity of the noise term *k*_*r*_ in the system (although one would expect tectonic movements to significantly change the short-term dynamics). Interestingly, this allows for the system to be “noisy”, i.e. many functional stem cells to contribute to the long-term dynamics, without any clonal dispersion, showing that one must be careful in equating the two directly. This would in particular be relevant for the dynamics of intestinal crypts, where cells away from starting position 0 have been shown experimentally to still contribute long-term (see Fig. 2 of the main text), but where little clonal fragmentation was observed (raising the possibility that such tectonic collective movements could occur to reposition cells towards/away from the best location).

## Appendix D: Numerical simulations and parameter estimation

### 1D simulation

A one-dimensional array of 20 cells was initialised, where every cell was given an index *x* = {0, 1, …, 19} to identify their starting position in the crypt, corresponding to their lineage. 0 is the most advantageous position at the bottom of the crypt, and 19 is at the top where it will be removed from the crypt by any single division event below. The simulation parameters were *k*_*d*_, the probability that a cell divides, and *k*_*r*_, the probability that a cell switches positions with its neighbour in either direction. During the simulation, cells were chosen at random with equal probability to decide whether to divide or change neighbors. The simulation was terminated when the array was colonized by a single cell lineage –we denote this as a “win” by that lineage.

The probability of survival of a given lineage, *p*(*c*_*k*_) was calculated as the number of wins divided by the number of simulation runs. The simulation was repeated 2000 times. For every 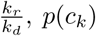 was plotted as a function of the starting position *k*, and the data was fitted with the following function:

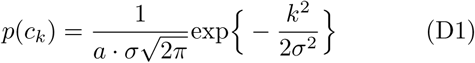

where *a* is a normalization constant. Consistently with the theoretical predictions given in equation (A13), we find that:

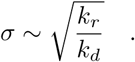

### 2D simulation

A two-dimensional array of size 8×20 cells was initialised, where every cell was given an index *x* = {0, 1, …, 19} to identify their starting row in the crypt, corresponding to their lineage. 0 is the most advantageous row and 20 is the upper boundary after which cells are removed (Note that this boundary condition is largely irrelevant for the results because cells at these rows have vanishing chance to contribute to a winning lineage). Periodic boundary conditions were applied to every row. This simulation uses the same parameters *k*_*d*_ and *k*_*r*_, but now with a maximum of 4 possible neighbours for intercalation. The simulation was repeated 2000 times, and plotting the probability of survival of a given lineage shows that it fits to:

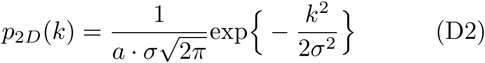

where *a* is a normalization constant. In agreement with the theoretical predictions given in equation (A13), we find that:

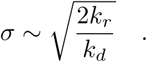

Both fits for intestinal homeostatic crypts and developing kidney shown in the main text were performed using this 2D simulation. In both cases, in order to build confidence intervals, in a non-parametric way, for our predictions of clonal survival as a function of position, we simulated 2000 times the number *N* of labelling events as in the data (*N* = 45 for intestine, *N* = 24 for kidney), and calculated the mean survival probability, as well as the 68% confidence interval around this prediction (i.e. 1 standard deviation around the mean). In both cases, we found that all of the experimental datapoint was contained within this interval. Finally, although on the long-term, survival probabilities converge towards a steady-state universal gaussian distribution, the live-imaging datasets were experimentally acquired on finite timescales, requiring simulations to examine the dynamics of clonal conversion. We thus state below for each organ the duration of the simulations *T* (rescaled by the cell division rate *k*_*d*_).

For the intestinal crypts, clones were initialized only at positions 0,1,2,3 and 4, to match with the experimental set-up where clones were only traced from Lgr5+ stem cells. Moreover, we defined a clone as “lost” in the system as soon as it didn’t have any cells in this compartment (positions 0-4), again to match the way that the experimental data was recorded in Ref. [13]. One should note that this assumption is expected to be largely irrelevant for our findings, given the results of Section 5: *k*_*r*_*/k*_*d*_ in this system is small enough that *N*_*s*_ < 5, so that the probability for clones to come back in the Lgr5+ region after having left it during the early phase of clonal competition is vanishingly small. Plots in the main figure for intestinal crypts used the following parameters: *k*_*r*_*/k*_*d*_ = 1 and a runtime of *Tk*_*d*_ = 0.5 (we note that the latter is relatively small, corresponding experimentally to a bit less than one full cell division in 3 days in Ref. [13], which could be linked to the method of intravital imaging (typical timescales reported in intestine are 1-2 days).

For the kidney tips, clones were initialized evenly in positions 0-10, which was also the definition for the compartment of clonal survival. Plots in the main figure used the following parameters: *k*_*r*_*/k*_*d*_ = 16 (see main text for details on this parameter estimation) and a runtime of *Tk*_*d*_ = 2 (which is the typical average number of divisions seen in the experimental dataset during the time course, see Ref. [24]). The experimental data of Ref. [24] assigns to cells a starting position on a 10×10 grid, and notes whether a clone still remains in the tip at the end of the observation period. As position (10, 10) was the edge of the tip, we calculated the euclidian distance of all coordinates (*i, j*) from position (10, 10), which is the starting distance reported in Fig. 3 of the main text. One should note that because of the longer runtime of experiments, predictions in kidney are much-closer to their steady-state universal form.

Finally, we also used these 2D simulations to fit the clonal fragmentation seen in mammary gland (where no long-term live-imaging was possible to follow clonal survival). To match experiments where cells were labelled in mouse of 3 weeks and collected at 5 weeks, and where the typical cell division rate is 16 hours, we used *Tk*_*d*_ = 20, although the predictions are largely insensitive to this timing. We then measured for each labelled cell the distance to the closest labelled cell, and built probability distributions for these nearest cell-cell distance. As expected, for low values of *k*_*r*_*/k*_*d*_, cells are always close neighbours, whereas the average cell-cell distance increased monotonously with *k*_*r*_*/k*_*d*_. In figure (10) of this Appendix we describe the process of inference of the *k*_*r*_*/k*_*d*_ parameter. In Fig. 4 of the main text we report the explicit probability distributions for the nearest cell-cell distance inferred from clonal dispersion observed during the development of the mammary gland.

## Appendix E: Experimental procedures

**FIG. 10:**
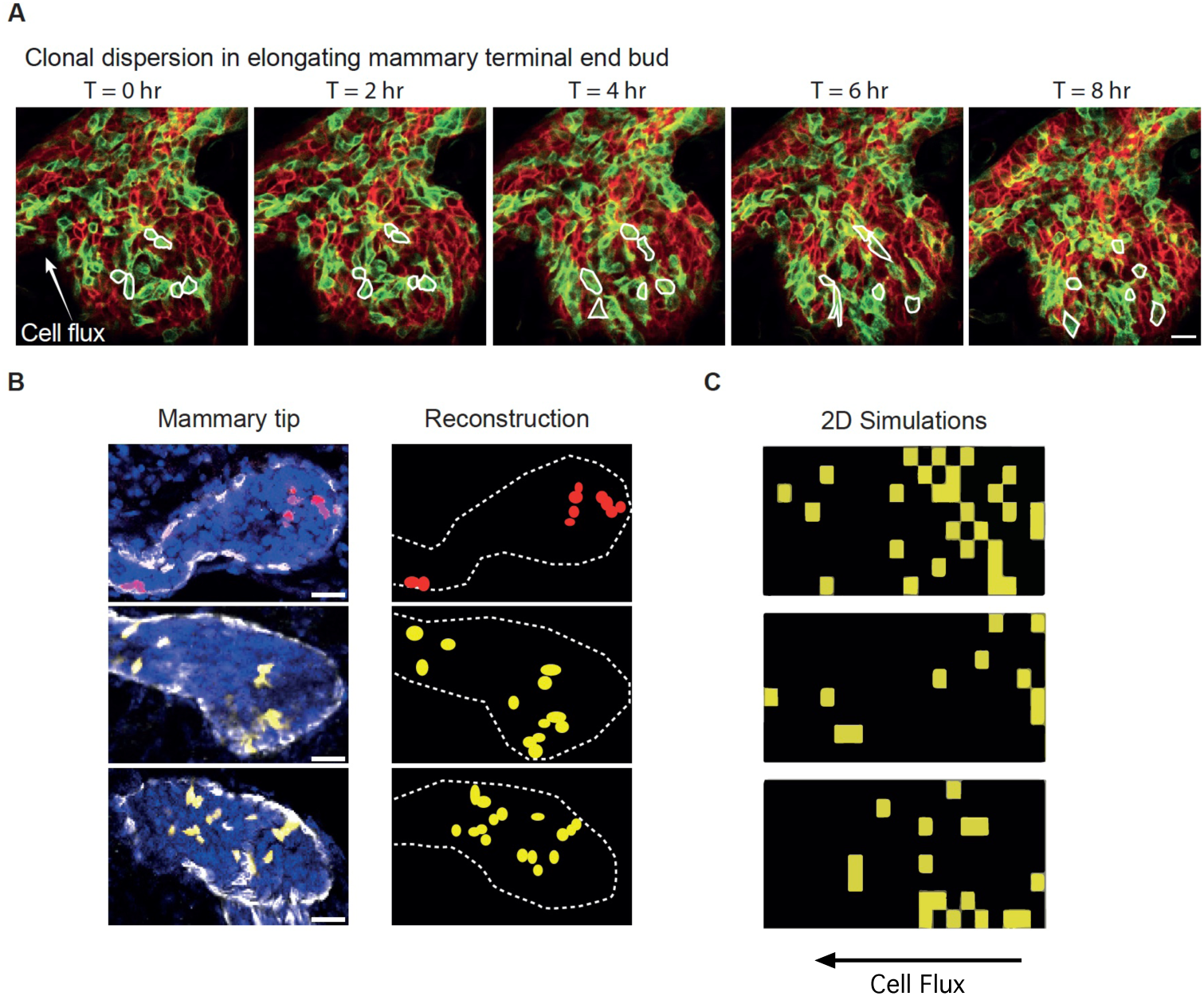
Inferring *k*_*r*_ from clone dispersion in mammary gland development A/ Intravital microscopy images of a developing terminal end bud followed over multiple hours showing extensive cell rearrangements leading to clonal dispersion. Some cells are highlighted with white lines to illustrate the random cell movements within the terminal end bud. Scale bar represents 10 *µ*m. B/ Confocal images of terminal end buds stained with keratin 14 (white) to label the basal cells, and DAP (blue) to label the nuclei. Cells from the same lineage are marked in red or yellow. The reconstruction of the mammary tips is shown in the right panel, from which the minimal distance between cells from the same lineage was inferred. Scale bar represents 25 *µ*m. C/ The real measures of the minimum distances obtained from direct observation of the clones are compared to simulations at different *k*_*r*_*/k*_*d*_’s, and the inferred *k*_*r*_*/k*_*d*_ for the real system is the one whose distribution of minimum distances is the closest to the real. In the pictures we show different snapshots corresponding to numerical simulations for the clone the fragmentation dynamics with *k*_*r*_*/k*_*d*_ = 3.

### 1. Mice

All mice were females from a mixed background, housed under standard laboratory conditions, and received food and water ad libitum. All experiments were performed in accordance with the Animal Welfare Committee of the Royal Netherlands Academy of Arts and Sciences, The Netherlands.

### 2. Intravital microscopy

R26-CreERT2;R26-mTmG female mice were IP injected with 0.2mg/25gTamoxifen diluted in sunflower oil (Sigma) at 3 weeks of age to induce sporadic recombination in the developing mammary gland. At 5 weeks of age, a mammary imaging window was implanted near the 4th and 5th mammary gland –for details, see [49]. Mice were anesthetized using isoflurane (1.5% isoflurane/medical air mixture) and placed in a facemask with a custom designed imaging box. Imaging was performed on an inverted Leica SP8 multiphoton microscope with a chameleon Vision-S (Coherent Inc., Santa Clare, CA, www.coherent.com), equipped with four HyD detectors: HyD1 (¡455nm), HyD2 (455–490nm), HyD3 (500–550nm) and HyD4 (560–650nm). Collagen I (second harmonic generation) was excited with a wavelength of 860nm and detected with HyD1, GFP and Tomato were excited with a wavelength of 960nm and detected with HyD3 and HyD4. Mammary gland tips were imaged at an interval of 20-30 minutes using a Z-step size of 3*µ*m over a minimum period of 8 hours. All images were acquired with a 25 × (HCX IRAPO N.A. 0.95 WD 2.5mm) water objective.

### 3. Quantitative image analysis

Clonal dispersion in the developing mammary tips was measured in lineage traced whole mount glands from R26-CreERt2;R26-Confetti mice as previously described [9]. In brief, R26-CreERt2;R26-Confetti female mice were injected at 3 weeks of age with 0.2mg/25g Tamoxifen to achieve clonal density labelling (¡1 cell per tip). Lineage traced mice were sacrificed at mid-puberty (5 weeks of age) or at the end of puberty (8 weeks). Mammary glands were dissected, fixed in periodate-lysing-paraformaldehyde (PLP) buffer (1% paraformaldehyde (PFA, Electron Microscopy Science), 0.01M sodium periodate, 0.075M L-lysine and 0.0375M P-buffer (0.081M Na2HPO4 and 0.019M NaH2PO4) (pH 7.4)) for 2 hours at room temperature (RT), and incubated for 2 hours in blocking buffer containing 1% bovine serum albumin (Roche Diagnostics), 5% normal goat serum (Monosan) and 0.8% Triton X-100 (Sigma-Aldrich) in PBS. Subsequently, glands were incubated with primary antibodies anti-K14 (rabbit, Covance, PRB155P, 1:700) or anti-E-cadherin (rat, eBioscience, 14-3249-82, 1:700), and secondary antibodies goat anti-rabbit or goat anti-rat, both conjugated to Alexa-647 (Life Technologies, A21245 and A21247 respectively, 1:400). Mammary glands were mounted on a microscopy slide with Vectashield hard set (H-1400, Vector Laboratories), and imaging was performed using a Leica TCS SP5 confocal microscope, equipped with a 405nm laser, an argon laser, a DPSS 561nm laser and a HeNe 633nm laser. All images were acquired with a 20x (HCX IRAPO N.A. 0.70 WD 0.5mm) dry objective using a Z-step size of 5*µ*m (total Z-stack around 200*µ*m). 3D tile scan images of whole-mount mammary glands were used to manually reconstruct the tips. Labelled confetti cells were annotated in the schematic outline of the tips including information on the confetti colour for the mammary glands (GFP=green, YFP=yellow, RPF=red and CFP=cyan). The length and the width of the tips were measured, and the co-ordinates of each labelled confetti cell in the tip were determined. The coordinates were used to calculate the position of each labelled cell within the ductal tip, as well as the minimal distance to the nearest neighboring cells with the same confetti color (as a measure of the clonal dispersion within the stem cell zone).

## Footnotes

1 A rigorous approach to this problem would require a reflecting boundary condition at *x* = 0. Imposing such boundary condition would make the whole problem much more difficult and, eventually intractable. The reason by which we adopted natural boundary conditions is due to the fact that the dynamics in this system is extremely imbalanced and runs essentially in only one direction. If one takes equation (A6) at *x* = 0 we observe that the probability of being at *x* = 0 decays as ∼ *e*^−*τ*^, for any starting point *k* > 0, as it is the case in our system. This tells us that the probability of visiting regions *x* < 0 is, to our purposes, negligible.

